# Agent-Guided De Novo Design of Nanobody Binders Against a Novel Cancer Target

**DOI:** 10.64898/2026.04.13.717816

**Authors:** Yue Zhao, Melih Yilmaz, Edward Lee, Chuanyui Teh, Lan Guo, Kemal Sonmez, Luca Giancardo, Gordon Trang, Fangda Xu, Madelyn Espinosa-Cotton, Nai-Kong V Cheung, Jiwon Kim, Xinyun (Nina) Cheng

## Abstract

Therapeutic antibody discovery remains slow and resource-intensive, with traditional methods providing limited control over epitope selection. We present a workflow for de novo nanobody design applied to a novel Desmoplastic Small Round Cell Tumor target encompassing four stages: (1) epitope identification guided by our hotspot recommendation agent using physical chemistry-based structure and sequence analysis tools with two curated databases (IEDB, PFAM), (2) de novo nanobody generation using three independent methods (RFantibody, IgGM, mBER) across multiple predicted antigen structures and nanobody frameworks, (3) multi-metric scoring including structural metrics from folding models, and *in silico* binding affinity from our sequence-based predictor, (4) high-throughput yeast surface display (YSD) screening followed by surface plasmon resonance (SPR) characterization of the specific binders. We generated 288,000 nanobody designs spanning eight target epitope regions and three variable domains of heavy chain-only antibody (VHH) frameworks. Multi-objective Pareto filtering with our candidate selection agent yielded 100,000 candidates for YSD screening with fluorescence-activated cell sorting (FACS). Of 116 enriched candidates advanced to SPR characterization, 46/116 (39.7%) produced reliable kinetic fits with *R*_max_ ≥ 30 RU, yielding *K_D_* values from 0.66 nM to 305 nM (median 31.7 nM). These results show that an agent-guided computational workflow can design nanomolar to sub-nanomolar nanobody binders against a novel target without experimental structure or prior antibody information.

## 1 Introduction

Monoclonal antibodies have become the dominant modality in biopharmaceutical development, with over 100 FDA-approved antibody drugs and well-established manufacturing, regulatory, and clinical development pathways [1]. Specialized formats such as single-domain VHHs (nanobodies) further expand therapeutic possibilities through their favorable developability characteristics, including high solubility, robust expression yields, low aggregation propensity, excellent thermal and chemical stability, and ease of engineering [2].

Yet conventional antibody discovery remains fundamentally constrained. Animal immunization, phage display, and synthetic library screening have proven success but impose inherent restrictions on accessible sequence space and demand months of iterative optimization for affinity, stability, and immunogenicity to ensure fitness for purpose [3, 4]. Most critically, these methods provide minimal prospective control over epitope selection, sequence composition, and paratope architecture, which can ultimately determine therapeutic success. Cryptic epitopes often prove intractable, and initial hit sets offer no guarantee of viable clinical candidates.

Machine learning has transformed our antibody predictive and design capabilities. Structure prediction methods including AlphaFold [5, 6], Boltz [7], Chai [8], and IntelliFold [9] now enable accurate modeling of protein structures, while specialized tools like NanobodyBuilder2 [10] predict VHH conformations with high confidence. For de novo antibody design, diffusion-based frameworks including RFdiffusion [11] and BindCraft [12] have successfully designed novel protein binders with explicit epitope control. Complementary approaches such as IgGM [13], which jointly optimizes complementarity-determining region (CDR) sequences and structures through diffusion processes, and mBER [14], which leverages backpropagation through structure prediction models, expand the design methodology toolkit. Sequence-based predictors utilizing protein language models like ESM2 [15] provide orthogonal validation of binding predictions independent of structural modeling assumptions. However, turning computational designs into functional antibodies presents distinct challenges: not only does generating these molecules require precise spatial positioning and conformational dynamics for effective target engagement, but integrating diverse machine learning models and tools into a cohesive discovery pipeline remains a bottleneck. Recent efforts including RFantibody [16] and Germinal [17] demonstrate modest experimental success rates, with reported hit rates ranging from 0.1% to 39% [16, 17, 18, 19], though direct comparison across studies is difficult because hit rate definitions vary (e.g., ELISA positivity vs. SPR-confirmed binding), targets differ in difficulty, and screening funnels differ in stringency.

In this study, we test these approaches against a target that amplifies these difficulties: a potential novel cancer target for Desmoplastic Small Round Cell Tumor, identified through transcriptomic analysis at Memorial Sloan Kettering Cancer Center (MSK) (see Methods: Target Identification for details). This target presents a series of challenges that test the generalization of current de novo antibody design. Without a wet-lab-resolved protein structure from cryo-electron microscopy or X-ray crystallography, epitope identification and interface modeling must rely entirely on computationally predicted structures, which—despite remarkable recent progress— can introduce uncertainties in surface-exposed loop conformations, side-chain orientations, and local electrostatic features critical for defining druggable binding sites. These uncertainties are compounded by the target’s length and structural complexity; the multi-domain architecture, flexible interdomain regions, and potential glycosylation sites are poorly captured by static predictions, making it difficult to confidently identify stable, accessible, and functionally relevant epitopes for nanobody engagement.

Also, no public antibody information exists for this target, eliminating conventional starting points such as paratope-epitope template transfer, homology-based CDR grafting, or affinity maturation trajectories from prior campaigns. The absence of any antigen-antibody interaction data for this target means that neither the de novo design models employed in this work, i.e., RFantibody, IgGM, or mBER, nor the structure prediction and binding affinity models used for computational filtering, have encountered this specific antigen target during training. However, these models do carry transferable representations of antibody-antigen binding principles learned from other complexes in the PDB. The challenge therefore lies in whether these generalized priors can produce functional binders against an antigen surface not seen during training. To date, most de novo antibody design campaigns have focused on well-characterized antigens with known structures and prior antibody data; efforts to validate these approaches against truly novel targets remain limited and have only recently emerged [20].

Here, we report the *in silico* and *in vitro* results from the first cycle of designing nanobodies against this target through an integrated computational-experimental workflow. Our workflow encompasses four stages: (1) generating target structures and identifying epitopes with favorable binding characteristics; (2) computationally designing 288,000 nanobody candidates constrained by epitope definitions while maintaining format-specific requirements; (3) computationally validating and filtering candidates at scale; (4) *in vitro* validation of 100,000 candidates binding to the target antigen using yeast display, FACS and SPR. This initial demonstration establishes that an integrated computational-experimental workflow can produce nanomolar to sub-nanomolar affinity nanobody binders against an uncharacterized target lacking public antibody sequence or experimental structures, providing a foundation for iterative optimization through subsequent design-build-test-learn (DBTL) cycles.

## 2 Results

### 2.1 De novo designed nanobodies bind a novel cancer target with nanomolar to sub-nanomolar affinity

Following the computational design and scoring pipeline (Figure 1), we identified epitope residues using our internally developed hotspot recommendation agent and de novo designed 288,000 nanobody candidates, which were filtered down to 100,000 using our candidate selection agent performing multi-property optimization. The 100,000 candidates were screened by yeast surface display (with a 90.6% VHH expression rate in yeast library) with 2 rounds of FACS enrichment where antigen-specific binders were identified by flow cytometry using individually stained yeast clones. 116 candidates were advanced to SPR characterization based on mean fluorescence intensity (MFI) at the 2nd round of FACS (Figure S1). 116 candidates expressed successfully, with a median yield of 184 mg/L (range: 34.5–200 mg/L) at the 2 mL culture scale, with 97% of the library exceeding 100 mg/L.

**Figure 1.**
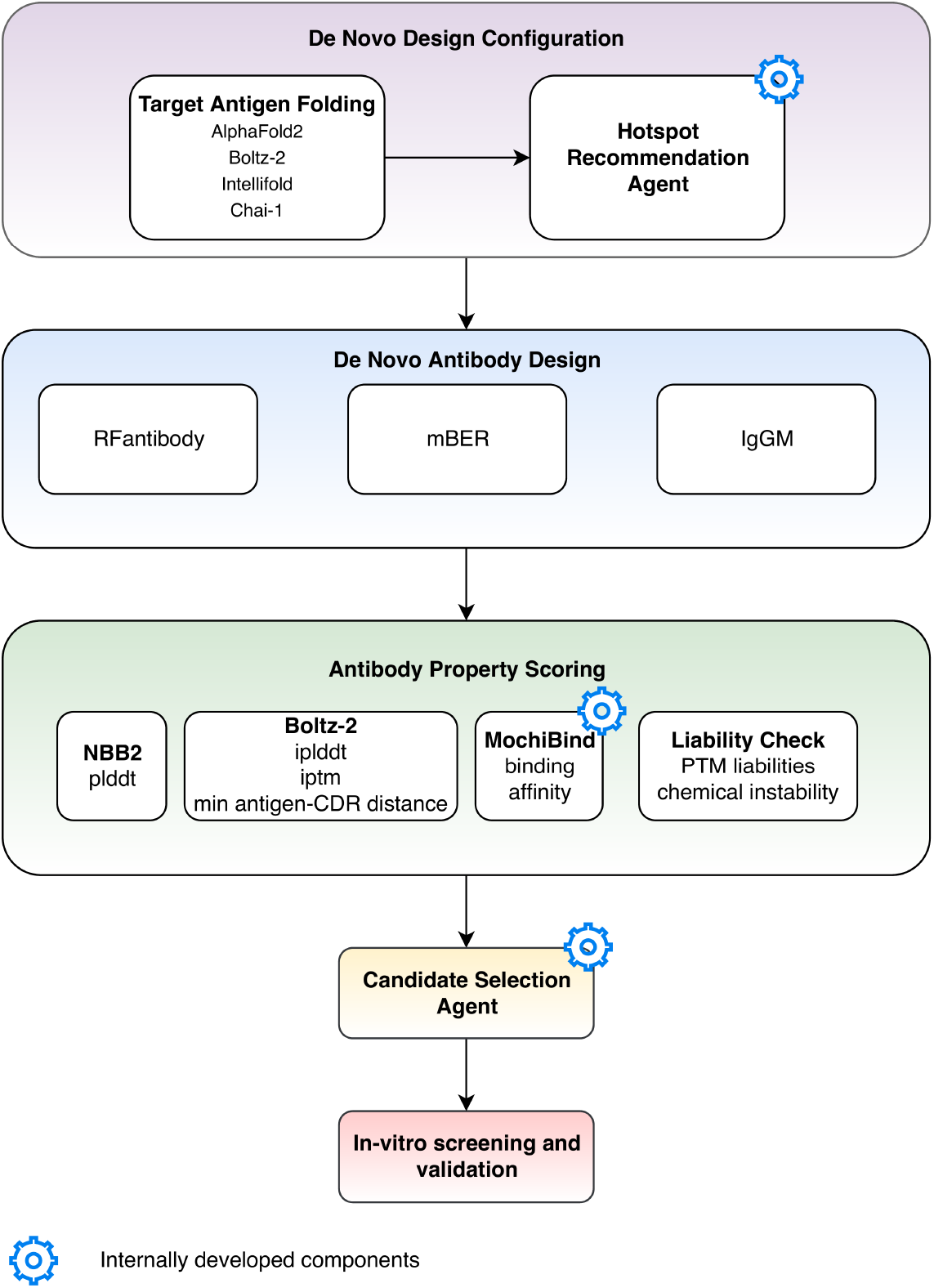
Integrated computational and experimental pipeline for de novo nanobody design. The workflow proceeds through five stages: (1) De Novo Design Configuration’s (purple) hotspot recommendation agent predicts target antigen structures using four folding methods and recommends 8 binding epitopes. (2) De novo design (blue) generates 288,000 nanobody candidates using three complementary methods (RFantibody, mBER, IgGM) across multiple epitopes and frameworks. (3) Multi-property scoring (green) evaluates all 288,000 candidates using structural metrics (NanobodyBuilder2 pLDDT, Boltz-2 complex quality and CDR-antigen distance), sequencebased binding affinity prediction (MochiBind [21], see Methods section for a more detailed description of this model), and antibody sequence liability checks. (4) Candidate Selection Agent (yellow) performs multi-objective optimization to select 100,000 top candidates. (5) *In vitro* screening and validation (pink) experimentally validates binding activity, with 116 candidates advanced to SPR characterization, yielding 46 confirmed binders (*R*_max_ ≥ 30 RU) with measurable kinetics. Gear icons denote internally developed components that automate key decision-making steps in the workflow.

Specificity testing confirmed no binding to an irrelevant antigen, transferrin receptor protein (TfR1), across all 116 candidates. Of the 116 candidates characterized, 46 (39.7%) produced binding responses to the target antigen with both measurable *K*_*D*_ and *R*_max_ ≥ 30 RU, providing reliable kinetic measurements. These 46 binders spanned *K*_*D*_ values from 0.66 nM to 305 nM, with a median *K*_*D*_ of 31.7 nM. The top candidate, PRJ266_044, achieved *K*_*D*_ = 0.66 nM. The second-ranked candidate, PRJ266_080, measured *K*_*D*_ = 2.3 nM (Figure 2). An additional 29 candidates shared detectable but low-amplitude SPR responses (*R*_max_ < 30 RU). Low *R*_max_ values can arise from multiple factors including low active concentration of the binder, partial binder immunoreactivity, and the reduced reliability of kinetic parameter estimation with low signals. The fitted *K*_*D*_ values for these candidates should therefore be considered provisional estimates requiring orthogonal confirmation. Among them, 14 candidates yielded *K*_*D*_ values in the sub-nanomolar range, including PRJ266_104 (*K*_*D*_ = 0.13 nM, *R*_max_ = 12.1 RU). SPR sensorgrams for the three candidates PRJ266_044, PRJ266_080, and PRJ266_104 are shown in Figure S2.

**Figure 2.**
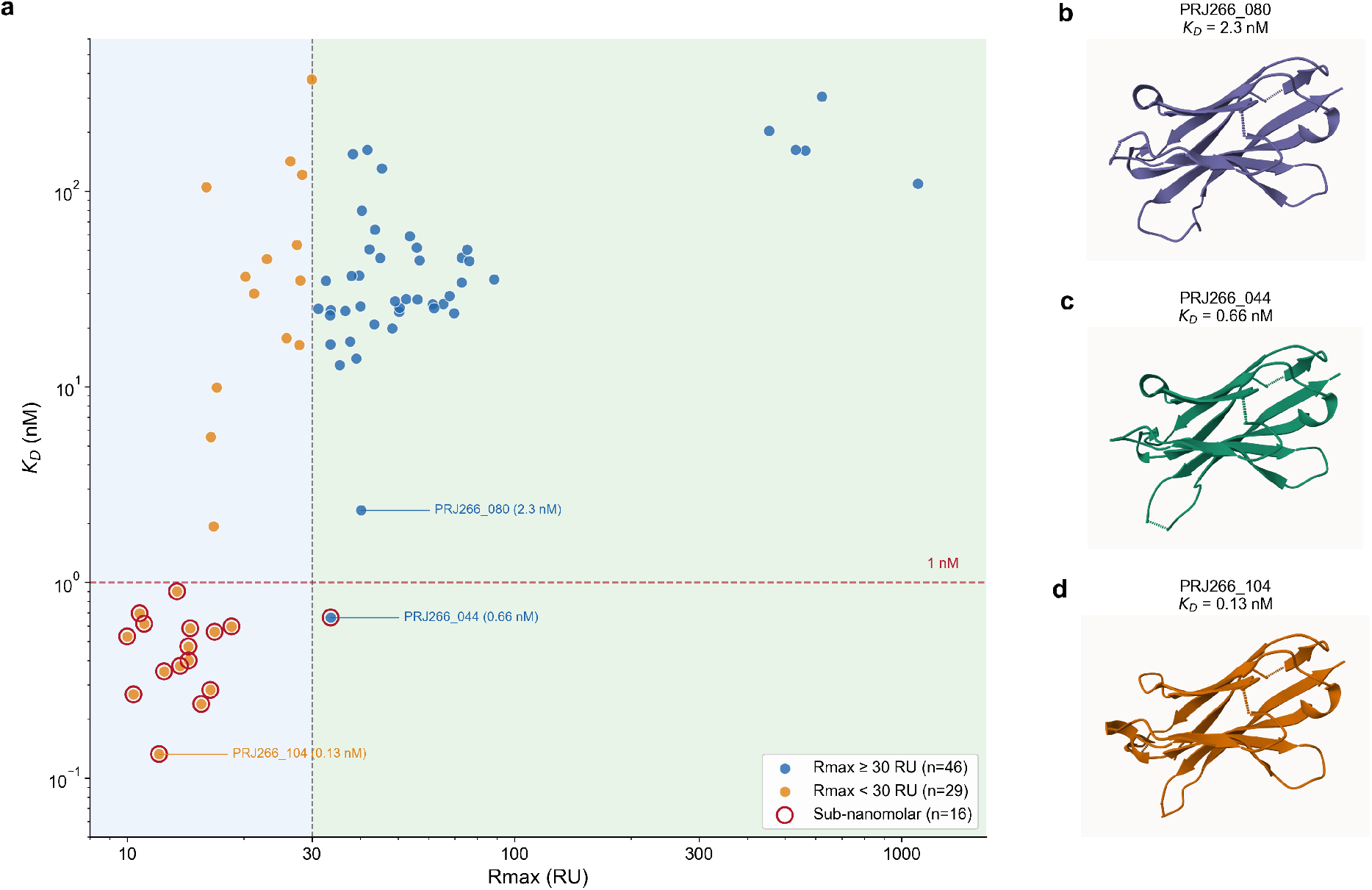
Binding affinity landscape and predicted structures of top de novo designed VHH antibodies. (a) Equilibrium dissociation constant (*K*_*D*_) plotted against SPR maximum response (*R*_max_) for all 75 candidates with measurable target binding. Blue points denote candidates with *R*_max_ ≥ 30 RU (n = 46), a threshold (dashed vertical line) above which kinetic fits benefit from favorable signal-to-noise; orange points denote candidates below this threshold (n = 29). Red open circles highlight 16 candidates with sub-nanomolar affinity (*K*_*D*_ < 1 nM; dashed horizontal line). Values for candidates with *R*_max_ < 30 RU carry substantially higher uncertainty due to limited signal amplitude and should be interpreted as provisional estimates. Three top-performing candidates are annotated: PRJ266_044 (*K*_*D*_ = 0.66 nM), the highest-affinity confirmed binder with *R*_max_ ≥ 30 RU; PRJ266_080 (*K*_*D*_ = 2.3 nM), the second-ranked binder in that group; and PRJ266_104 (*K*_*D*_ = 0.13 nM), the lowest measured *K*_*D*_ overall, though this estimate carries substantial uncertainty due to low *R*_max_. (b–d) Predicted structures of the three annotated candidates, generated using NanobodyBuilder2.

We examined the influence of design parameters on binding affinity among the 46 high-signal binders (Figure 3). Of the three nanobody frameworks used in our de novo design campaign, only two yielded confirmed binders in the high-signal group (*R*_max_ ≥ 30 RU), and the distribution was heavily skewed: Framework B accounted for 45 of the 46 high-signal binders (Figure 3a). This dominance indicates that framework properties can determine whether computationally designed sequences translate into strong binders and highlights the value of diversifying frameworks as a design parameter to avoid missing productive scaffolds. Both IgGM and mBER produced binders in the *R*_max_ ≥ 30 group, with IgGM contributing the larger share at a lower median *K*_*D*_ (n = 33, median 28.0 nM) compared to mBER (n = 13, median 43.9 nM), though this difference was not statistically significant (p = 0.311) (Figure 3b). All three RFantibody-designed binders returned fitted *K*_*D*_ values in the sub-nanomolar to low-nanomolar range (0.13, 0.62, and 5.5 nM) but with uniformly low *R*_max_ values (11.1-16.4 RU), all below the 30 RU threshold. The low SPR signal increases uncertainty in the kinetic fits, so these affinity estimates carry wider confidence intervals than those from higher-*R*_max_ candidates. Across the five structure prediction methods used to generate input antigen conformations: Boltz-2 (with and without potential-guided variant), IntelliFold, Chai-1, and AlphaFold2, no statistically significant difference in median affinities was detected (26.5-47.5 nM, Kruskal-Wallis p = 0.330, Figure 3c). However, with only 46 binders split across five groups (n = 3-13 per group), this analysis is substantially underpowered, and meaningful differences in affinity across folding models cannot be ruled out.

**Figure 3.**
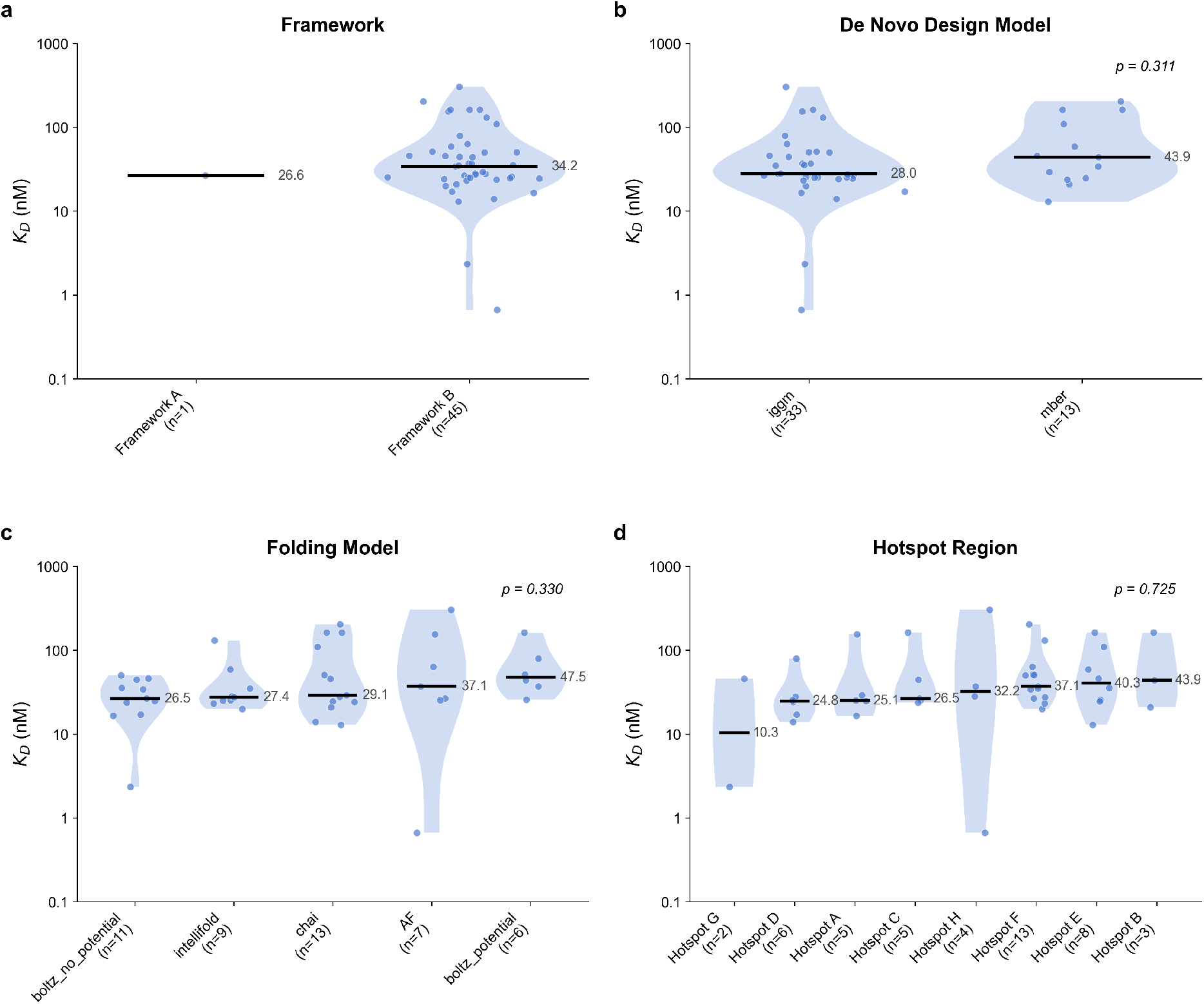
Binding affinity distribution by design parameters. Violin and strip plots of *K*_*D*_ for 46 confirmed binders (*R*_max_ ≥ 30 RU) grouped by (a) structural framework, (b) de novo generative model, (c) structure prediction model, and (d) target hotspot region. Black lines indicate group medians (nM). p-values from Kruskal–Wallis test.

Binders originating from all eight hotspot-conditioned design groups were confirmed by SPR, with median *K*_*D*_ values ranging from 10.3 nM (Hotspot G) to 43.9 nM (Hotspot B) (Figure 3d), though the small per-group sample sizes preclude meaningful comparison across groups (Kruskal-Wallis p = 0.725). Boltz-2 co-folding of full-length antigen and antibody sequences independently confirms that the designs concentrate contacts near agent-recommended hotspots (Figure 4b), but the mapping between conditioning hotspot and predicted binding site is not strictly one-to-one. A sequence conditioned on one hotspot may dock at a neighboring or overlapping hotspot in the co-folded structure, particularly where hotspots share a spatially contiguous surface. In practice, this cross-hotspot engagement is not detrimental to the workflow: because the agent identified all eight regions as viable binding sites prior to design, a sequence that engages any agent-recommended hotspot (whether or not it matches its conditioning target) still represents a productive design outcome. SPR confirms target binding but does not resolve which epitope is engaged; experimental epitope mapping would be needed to determine the true binding sites.

**Figure 4.**
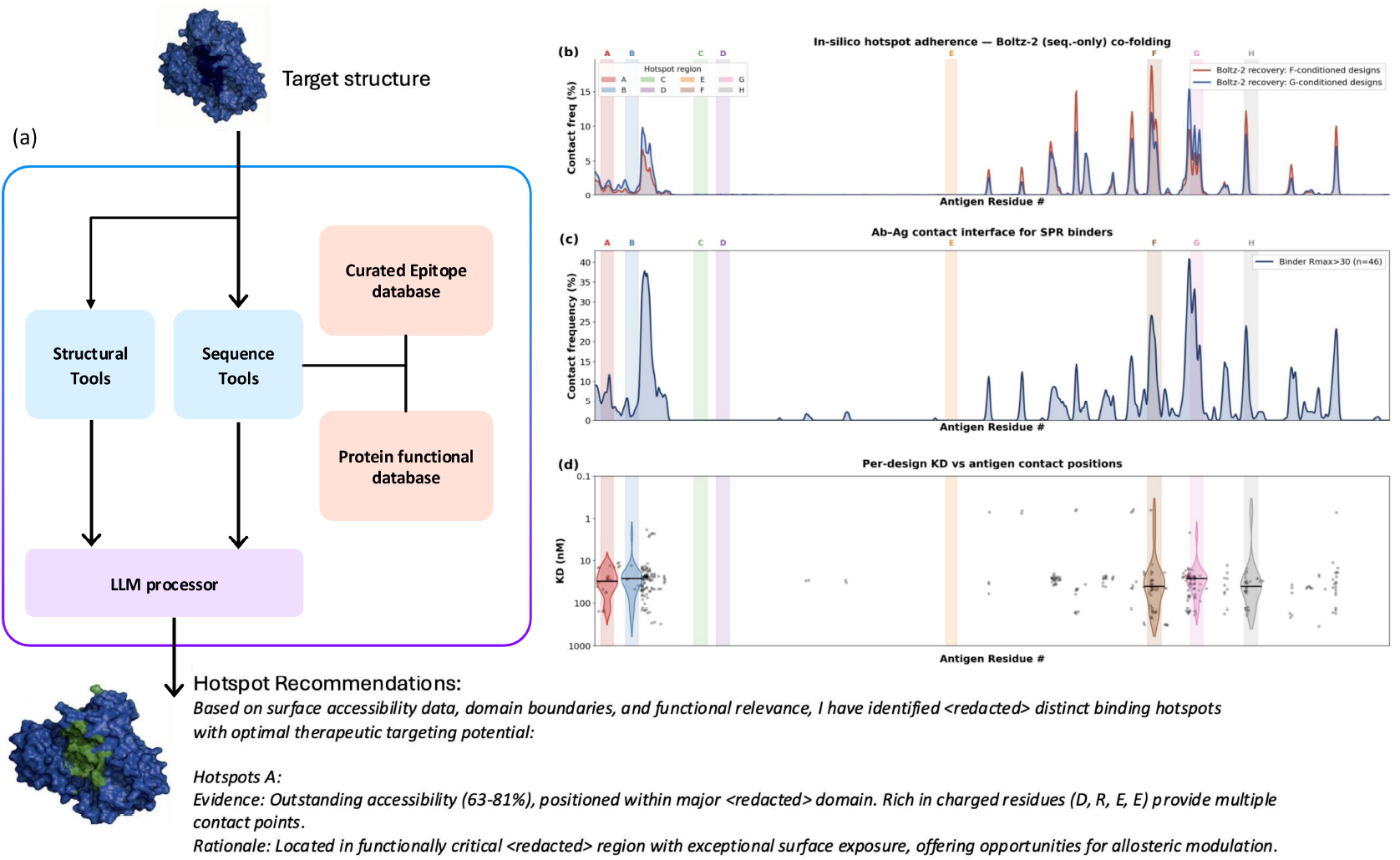
Hotspot Recommendation agent. (a) The agent analyzes the structures of the target (PDB) and recommends hotspot residues that are used to condition binder design to the target. (b) *In silico* hotspot adherence. The recommended hotspots are supplied as input to de novo structure-based design models (RFantibody, IgGM, mBER) which generate VHH sequences conditioned on each hotspot. The designed sequences are then co-folded with the antigen using Boltz-2, a sequence-only structure prediction model without previous hotspot and structural knowledge. The overlay shows that Boltz-2 recovers the hotspot-conditioned interface: designs targeting the agent’s proposed hotspots engage near the intended targets. Nonetheless, we observe epitopes that are independent of hotspot conditioning as some epitopes are highly surface accessible and energetically favorable. (c) Antibody– antigen contact interface after Boltz-2 complex folding showing CDR contact frequency (% of binder designs contacting the i-th antigen residue) for 46 SPR-confirmed binders (*K*_*D*_ measurable, *R*_max_ ≥ 30 RU). Contacts are defined as any CDR heavy atom within 7 Å of any antigen heavy atom in Boltz-2 predicted structures. Shaded vertical bands mark the 8 agent-defined hotspot regions. (d) Per-design *K*_*D*_ vs antigen contact positions. Each design has a single *K*_*D*_ value (y-axis) but many designs contact multiple antigen residues simultaneously; each contacted residue is plotted as a dot at the design’s *K*_*D*_ . Density estimates in (d) are shown for binders whose contacts fall within ±10 residues of each hotspot range.

### 2.2 Target Structure Prediction

To assess inter-method agreement, we computed pairwise TM-scores (Template Modeling score) among the five predicted structures after C*α* superposition (Table S1). TM-score is a metric for measuring structural similarity between protein models, ranging from 0 to 1, where scores >0.5 generally indicate proteins share the same overall fold [22]. TM-scores were computed on C*α* atoms after optimal superposition using TM-align. All pairs exceeded a TM-score of 0.5, confirming that every method recovered the same overall fold. The four approaches: Boltz-2 (with and without potential-guided variant), Chai-1, and IntelliFold formed a tight cluster (TM-scores 0.78-0.91), with Boltz-2 and its potential-guided variant showing the highest agreement (TM = 0.91). Chai-1 and IntelliFold also converged closely (TM = 0.89). AlphaFold2 was consistently the most divergent prediction (TM = 0.62-0.66 against all other methods). The AlphaFold2 structure was retrieved from the AlphaFold Protein Structure Database [23] rather than generated with the same controlled parameters used for other structures, so differences in MSA construction, model version, or template usage may contribute to this divergence. The observed difference may therefore reflect both algorithmic and parametric factors.

### 2.3 De Novo Design – Hotspot recommendation

Designing nanobodies against a novel target with no prior structural or existing antibody-antigen interface data requires identifying suitable epitopes. To systematically identify candidate binding hotspots on the target antigen, we developed an AI agent that orchestrates multiple bioinformatics tools and synthesizes their outputs through the large language model (LLM), Claude Sonnet 4. The agent integrates a diverse set of tools operating on both the structure and sequence of the target protein, including retrieval of known epitopes and functional domains from two curated databases, to recommend hotspot residues for downstream de novo design (Figure 4a).

These tool outputs are provided as context to the LLM, which synthesizes the evidence to identify candidate binding hotspots. For each region, the LLM weighs multiple factors: the structural and sequence analyses from the tools, sequence divergence from user-specified negative controls, and any existing antibody binding sites in the target structure when the structure is a complex. By grounding the LLM’s recommendations in deterministic, verifiable tool outputs— computed surface accessibility, IEDB alignment hits, PFAM domain boundaries—rather than relying on the model’s parametric knowledge of protein biology, this architecture constrains the recommendation space to evidence-supported regions and reduces the risk of hallucinated epitope suggestions. The agent’s recommendation strategy prioritizes broad epitope coverage over narrow precision. In early-stage de novo design against an uncharacterized target, the cost of missing a viable epitope outweighs the cost of testing additional candidates.

Each of the agent’s bioinformatics tools is deterministic and individually validated. However, integration into the agent workflow required additional tool-level validation to ensure correct behavior on diverse PDB inputs. For structure-based tools, we verified that analyses are performed on the true antigen surface: when a researcher submits an antibody-antigen complex PDB, existing antibody chains must be removed before computing surface accessibility, secondary structure, and hydrophobicity, as the presence of artificial binding partners distorts the solvent-accessible surface. For database tools such as IEDB, MMseqs2 alignment parameters were tuned to maximize epitope recovery: default E-value thresholds penalize short sequences, yet most antibody epitopes are discontinuous and composed of smaller linear B-cell epitope fragments, requiring relaxed thresholds to avoid false negatives.

The end-to-end hotspot prediction capability was validated on antibody-antigen complexes from the Structural Antibody Database (SAbDab) [24]. A test set (n=320) was selected for sequence dissimilarity using clustering at 70% identity, and a curated holdout set (n=76) was constructed from immunologically relevant targets including viral antigens, tumor-associated antigens, and therapeutic targets. Epitope recovery was measured using a top-k prediction framework: the agent proposes k=5 non-overlapping 10-amino-acid regions per antigen, and a prediction is scored as a hit if any proposed region overlaps with at least one true epitope residue from the experimental complex structure. On the diverse holdout set, the agent achieved ∼80% top-5 accuracy.

### 2.4 De Novo Design – VHH binder generation

Our de novo design strategy employs three generative models: RFantibody, IgGM, and mBER, each representing a distinct computational approach to antibody design: diffusion-based backbone generation (RFantibody), joint sequence-structure diffusion (IgGM), and backpropagation-guided sequence optimization through structure prediction (mBER), respectively. By leveraging methods with fundamentally different design principles, we aimed to maximize sequence and structural diversity in the resulting candidate pool while reducing the risk of shared failure modes from any single approach. To maximize coverage of the design space, we varied multiple design parameters in a combinatorial fashion: eight computationally identified epitope hotspots, three distinct nanobody frameworks, and CDR III loop lengths ranging from 4 to 13 amino acids. Each combination was sampled across all three generative models, ensuring broad exploration across epitope, scaffold, and structural dimensions. To account for structural uncertainty of the target antigen, for which no experimental structure exists, we generated predictions using four structure prediction models: AlphaFold2, Boltz-2, Chai-1, and IntelliFold. These models were selected as leading structure prediction methods employing distinct architectural and algorithmic approaches, including evoformer-based prediction (AlphaFold2) [6], diffusion-based generation (Boltz-2) [7, 25], and alternative learned energy frameworks (Chai-1, IntelliFold) [8, 26], thereby maximizing structural diversity in the predicted conformations. Designing against multiple predicted conformations reduces the risk of overfitting to artifacts of any single model and increases the likelihood that generated candidates are robust to conformational variability of the true target structure. This multi-dimensional exploration strategy generated a library of 288,000 nanobody candidates (Figure 1).

### 2.5 Hotspot adherence using Boltz-2 to predict complex structure

The designed antibody-antigen interface can only be assessed after independent structure prediction. The de novo backbone generation stage produces structural coordinates, but these represent idealized geometries conditioned on input constraints (hotspots) and antigen structure. To assess whether the designed sequences encode hotspot preference recoverable by an independent structure prediction method, we performed Boltz-2 co-folding, which predicts complex structures from antibody and antigen sequences alone without access to the original hotspot constraints. We note that Boltz-2 and the design models share training data from the PDB, so this analysis tests whether hotspot specificity is encoded in the designed sequences and recoverable by co-folding, rather than providing fully independent validation of binding site accuracy. Interface quality was evaluated by computing pairwise distances between heavy atoms of heavy chain CDR side chains and heavy atoms of the target side chains.

The recommended hotspots were supplied as input to de novo structure-based design models (RFantibody, IgGM, mBER), which generated VHH sequences conditioned on each hotspot. The designed sequences were then co-folded with the antigen using independent structure prediction using Boltz-2, a sequence-only structure prediction model. Figure 4b illustrates the contacts of the interface as predicted by Boltz-2. Designs targeting the agent-recommended hotspots concentrate near the intended residues. This suggests that epitope specificity encoded during backbone generation is preserved through independent structure prediction, though both the design models and Boltz-2 were trained on overlapping PDB data, so shared biases cannot be excluded.

In Figure 4b, designs targeting Hotspots F, G, and H achieve high peak CDR contact frequencies at the intended epitope, and a clear distribution shift distinguishes designs conditioned on Hotspot F. These results confirm that the agent’s hotspot recommendations translate into distinct, recoverable binding site preferences under independent structure prediction. Notably, all profiles share a baseline of hotspot-independent contacts, particularly at the N-terminal region of the antigen. The agent recommends Hotspot B as it exhibits up to 90% surface accessibility and is energetically favorable with hydrophobic residues. Structural analysis of the candidates suggests Hotspot B forms part of a discontinuous epitope shared with Hotspots F and G, which would account for the overlapping contact signals observed across these regions.

Figure 4c shows contact frequency profiles for SPR-confirmed binders. After aggregating across all hotspots, the post-SPR filtering contact landscape broadly mirrors the pre-filtering distribution. Modest shifts are observed upon filtering: designs targeting Hotspots B, F and G appear enriched among the lowest-*K*_*D*_ binders, though this observation is based on small per-group sample sizes and should be treated as hypothesis-generating rather than conclusive. Combined with the Boltz-2 recovery of conditioned epitopes *in silico*, these results suggest that designs conditioned on Hotspots B, F, and G may engage a shared or overlapping binding surface. However, the SPR data confirms binding but does not determine the epitope. Experimental epitope mapping is needed to validate whether the computationally predicted binding sites correspond to the true epitopes.

### 2.6 *In Silico* Scoring, Filtering, and Candidate Selection of Diverse Antibodies

To prioritize candidates for experimental validation, we developed an *in silico* scoring and filtering framework combining structural, sequence-based, and developability metrics, followed by automated candidate selection through our candidate selection agent. Structural quality was assessed through antibody folding pLDDT scores, while complex formation and binding geometry were evaluated using three metrics from predicted antibody-antigen complexes: interface predicted template modeling score (ipTM), complex ipLDDT, and minimum CDR-antigen distance. To provide orthogonal validation independent of structural modeling assumptions, we applied MochiBind, a sequence-based binding affinity predictor developed by AWS Life Sciences (manuscript currently under peer review). MochiBind encodes antibody and antigen sequences using a pretrained protein language model (ESM-2) and predicts relative binding affinity through pairwise learning, without requiring structural information. We additionally performed developability assessments including systematic liability checks adapted from the Liability Antibody Profiling (LAP) methodology [27], which screen for sequence motifs associated with high-risk modifications such as N-linked glycosylation, deamidation, fragmentation, and oxidation within CDR and framework regions (see details in Methods: Antibody Diversity and Liability Analysis). These complementary metrics were integrated by our candidate selection agent, which employs multi-objective Pareto optimization to intelligently balance competing design objectives. Rather than collapsing quality into a single weighted score, which would require arbitrary assumptions about relative importance, the agent stratifies candidates into successive Pareto fronts, preserving diverse high-quality designs that excel across different metric combinations. This approach identifies non-dominated antibody ranks where improving any single property necessarily trades off against others, enabling systematic exploration of the design space while maintaining interpretable selection criteria (see Methods: Candidate Selection for details). Working in tandem with the hotspot recommendation agent that defined binding targets, the candidate selection agent automated the critical decision of which designs to advance, reducing 288,000 candidates to 100,000 sequences for experimental validation based on rigorous multi-property optimization rather than manual curation (see metrics distribution in Figure S3).

Our analysis revealed that the three-method design approach successfully generated diverse candidates across all design parameters. RFantibody, IgGM, and mBER produced non-overlapping sequence sets, confirming that methods with distinct design principles explore different regions of sequence space. To understand how design parameters affect the quality of designed antibodies, we analyzed the distribution of *in silico* scoring metrics across four design dimensions: target epitope, nanobody framework, seed antigen structure, and de novo design method. Among these, the choice of de novo method produced the most pronounced differences: RFantibody-designed nanobodies showed lower median scores for complex ipTM and CDR-antigen proximity compared to mBER and IgGM, while performing comparably on monomer folding quality and predicted binding affinity (Figure S3). Framework-dependent effects were also observed, with certain scaffolds consistently outperforming others in predicted binding metrics.

Sequence diversity analysis of the shortlisted candidates confirmed broadly distributed amino acid usage across interior CDR H1, H2, H3 positions, e.g. with longer CDR H3 loops (10-13 amino acids) exhibiting greater positional diversity than shorter loops, suggesting that design diversity naturally increases with loop length (Figure S4, S5 and S6 for CDR H1, H2, and H3 positional amino acid distribution analysis).

## 3 Discussion

Traditional antibody discovery methods, including animal immunization, phage display, and hybridoma technology, have a strong track record of success but suffer from significant temporal and resource constraints. A typical discovery campaign requires 6–12 months [28] from target identification to lead candidate selection, with animal immunization alone consuming 8–12 weeks [29] before any screening begins. Phage display libraries, while offering broader sequence diversity than immunization, still require multiple rounds of bio-panning (typically 3–5 rounds over 4–8 weeks [30]) followed by empiric affinity maturation campaigns involving iterative mutagenesis and reselection, which can substantially extend discovery timelines [31, 32]. Critically, these approaches provide minimal prospective control over epitope selection—the binding site is determined post hoc through structural characterization rather than being defined in advance. In contrast, our integrated *in silico* workflow generated 288,000 computationally diverse candidates with explicit epitope targeting across eight distinct binding regions. Where traditional methods screen 10^7^-10^9^ variants through a single biological selection, our approach systematically explored design space using three complementary generative methods (RFantibody [16], IgGM [13], mBER [14]) conditioned on agent-predicted hotspots, multiple target conformations, and varied frameworks. The resulting 100,000 experimentally validated candidates represent a scale of epitope-controlled diversity unattainable through conventional library construction, while the computational prefiltering based on complementary structure prediction (Boltz-2 [7]) and sequence-based binding affinity estimation enabled resource-efficient experimental validation. This approach shifts the discovery bottleneck from experimental screening throughput to the quality of generative models and scoring functions that provide computational metrics used to evaluate candidate antibodies, including structural predictions, sequence-based binding affinity estimates, and developability assessments.

The Hotspot recommendation agent considers both biophysical and biochemical accessibility to binding as well as antibody function. The agent synthesized inputs from IEDB, PFAM, surface accessibility, secondary structure, and hydrophobicity profiles to propose eight distinct hotspot regions, each supported by complementary lines of evidence. For example, Hotspot B was selected for its high surface accessibility (up to 90%) and cysteine content, while Hotspot G was prioritized based on its location within a functionally critical domain with favorable charged-residue composition. Crucially, these recommendations are not opaque rankings; the agent provides biophysical and functional rationale for each hotspot, grounding its selections in interpretable biophysical properties such as surface exposure, electrostatic favorability, and domain relevance. By narrowing the design space to structurally and functionally motivated epitopes, the agent provided a rational basis for de novo design against a target with no prior antibody data. The subsequent hotspot adherence analysis (Figure 4b-c) confirms that agent-recommended hotspots are recoverable by independent structure prediction, and SPR provided confirmation of binding.

Using data from both the adherence study and SPR data, the eight hotspots do not appear to function as fully independent epitopes. Hotspot B, despite being specified as a distinct region, shares overlapping contact signals with Hotspots F and G in the Boltz-2 co-folding analysis, suggesting these three regions have formed a spatially contiguous binding surface on the folded antigen. Among the confirmed binders, designs contacting a potential discontinuous epitope (B+G) achieved some of the lowest *K*_*D*_ values in the panel (e.g. candidate PRJ266_080 conditioned on hotspot G, 2.38 nM). This combinatorial pattern suggests that the agent’s individual hotspot recommendations, when viewed collectively, can reveal higher-order epitope architectures that no single hotspot definition would capture.

Filtering de novo candidates based on *in silico* metrics remains a challenge and when comparing the efficacy of various de novo methods and other parameters such as epitope and framework selection, focusing on the performance of median designs versus top percentile can yield different results. In our study, the choice of de novo method led to the most variance in terms of *in silico* metric on average, where RFantibody designed nanobodies were underrepresented in the filtered group, but the top designs had a balanced distribution across three de novo methods used in the study. We hypothesize this discrepancy arises from three factors: (1) architectural differences between diffusion-based backbone generation (RFantibody) and sequence-conditioned approaches (IgGM, mBER) may produce distinct score distributions even when generating high-quality binders, RFantibody’s structure-first paradigm optimizes geometric complementarity before sequence assignment, potentially yielding lower median sequence-based scores (e.g., MochiBind) while still producing top-percentile candidates with good binding geometry; (2) a known index offset issue in RFantibody’s hotspot conditioning may have introduced systematic positional errors that disproportionately affected median designs while top candidates overcame this through superior overall geometry; and (3) our use of both vanilla ProteinMPNN [33] and AbMPNN [34] for sequence design within the RFantibody pipeline may have contributed to score variance, as preliminary analysis of our *in silico* data suggests ProteinMPNN-designed sequences showed lower median binding affinity predictions compared to AbMPNN-designed sequences, though both methods produced comparable top-percentile candidates.

While *in silico* metrics enabled efficient filtering for library generation, we observed little correlation between most *in silico* folding metrics (pLDDT, ipTM) and experimental binding outcomes in the SPR data (Figure S7). This highlights that current structure prediction confidence scores, while valuable for assessing model quality, do not directly translate to robust binding affinity predictions, particularly for novel target antigens like the one used in this study, as observed by others in the literature, including mBER paper [14] where an increased ipTM filtering threshold for de novo designed nanobodies doesn’t translate to a higher experimental hit rate for various target antigens. This underscores the continued necessity of experimental validation at scale and points to opportunities for developing dedicated binding affinity predictors in future work.

Our approach has several limitations that we are actively working to address: (1) Limited assessment of biophysical properties, current reliance on *in silico* metrics (e.g. Boltz-2 co-folding, MochiBind affinity estimates) may not fully capture critical characteristics such as conformational stability or dynamic instabilities; we are developing molecular dynamics-based refinement to validate structural stability. (2) Limited scaffold diversity, the workflow used only three generative methods (RFantibody, IgGM, mBER), potentially constraining exploration of underrepresented scaffold classes that might offer superior developability; we are expanding scaffold diversity through additional generative models.

To integrate our *in silico* workflow more closely with experimental outcomes, we have designed a lab-in-the-loop framework [35] that will progress through Design-Build-Test-Learn (DBTL) cycles (Figure S8). The Design, Build and Test phases of the first cycle have been completed: computationally designed candidates were synthesized, displayed on yeast, and screened for binding activity, with binding-positive populations sequenced via NGS to link design parameters with experimental outcomes. From the initial screen, 116 candidates have been characterized by SPR for binding kinetics and affinity. Cell-based validation with target-expressing cell lines is planned to further characterize these candidates, followed by comprehensive developability characterization including thermal stability and polyreactivity to establish ground-truth fitness values. With this experimental foundation in place, the Learn phase remains: characterized binders will serve as training data to develop target-specific machine learning models for fitness prediction, with separate models for distinct properties (e.g., binding affinity, stability), which will then guide efficient directed evolution campaigns [36, 37, 38, 39, 40] where ML-predicted candidates are synthesized and experimentally validated, initiating subsequent DBTL cycles.

## 4 Conclusions

We designed nanobody binders de novo against a novel cancer target with neither an experimental structure nor prior antibody information. Starting from sequence alone, the workflow produced predicted target structures using four folding methods, identified eight candidate epitope regions through an agent that integrates physical chemistry-based analysis and curated database evidence (IEDB, PFAM), and generated 288,000 nanobody designs across three generative methods (RFantibody, IgGM, mBER), five predicted antigen conformations, and three VHH frameworks. Multi-metric scoring and Pareto-based filtering selected 100,000 candidates for yeast surface display screening. Of 116 candidates characterized by SPR, 46 produced reliable kinetic measurements (*R*_max_ ≥ 30 RU) with *K*_*D*_ values from 0.66 nM to 305 nM (median 31.7 nM). Both IgGM and mBER contributed confirmed binders in this group, and binders originating from all eight hotspot-conditioned design groups were represented among the confirmed hits, though SPR confirms target binding rather than engagement at the intended epitope. Specificity testing confirmed target-specific binding, with no candidates showing detectable interaction with the off-target control antigen.

The hotspot recommendation provided the epitope definitions used to condition all downstream design. Without prior antibody-antigen interaction data or experimental structures, conventional epitope selection strategies such as template transfer or homology-based CDR grafting were unavailable. The agent synthesized structural and sequence-level evidence, including surface accessibility, secondary structure, hydrophobicity, functional domain annotations, and known epitope databases, to propose eight binding regions, each supported by an explicit physical and functional rationale. Binders were recovered from designs conditioned on each of the eight hotspot regions. However, the true binding interfaces do not necessarily correspond to the conditioning hotspot. Boltz-2 co-folding analysis showed that designs conditioned on specific hotspots concentrated predicted contacts near the intended residues, and the lowest-*K*_*D*_ binders were enriched among designs targeting Hotspots B, F, G, and H. Experimental epitope mapping will be needed to confirm whether these predicted binding sites correspond to the true epitopes. Developability characterization, including thermal stability and polyreactivity has not yet been performed and is planned for subsequent work. The confirmed binders and the binder/non-binder labels derived from NGS sequencing of yeast display populations as well as the developability characterization will provide training data for target-specific ML models to guide the next design cycle, with the goal of improving binding affinity and developability through iterative optimization.

## 5 Methods

### 5.1 Target Identification

RNA sequencing was performed on 47 desmoplastic small round cell tumor (DSRCT) specimens collected from patients undergoing surgery at Memorial Sloan Kettering Cancer Center. All patients provided consent for their tumors to be used in research. We sought to identify highly expressed cell-surface targets for antibody-based therapy that are restricted in normal tissues to minimize on-target, off-tumor effects. To this end, we performed differential gene expression analysis to identify genes upregulated in our tumor samples compared to five normal tissue samples taken from the peritoneum of healthy donors. We narrowed this list to include only genes upregulated by EWSR1::WT1, the pathognomonic transcription factor present in all DSRCT and absent in normal tissue, and then to targets predicted to be expressed on the cell surface according to the *in silico* human surfaceome [41]. Finally, we excluded any genes with more than minimal expression in any normal tissue according to the Genotype-Tissue Expression (GTEx) database [42]. The target we selected was the most differentially upregulated gene remaining after these filters were applied to our dataset.

### 5.2 Stacks and Compute

To systematically generate and evaluate nanobody designs at scale, we implemented a batch processing framework using AWS HealthOmics [43]. A master manifest was created to orchestrate the design generation process, where each manifest entry specified a unique combination of design parameters including framework selection, target structure, hotspot definition, and other relevant variables. Each entry in the manifest corresponds to an individual Omics workflow execution that generated a predetermined number of designs, with the collective output across all runs totaling 96,000 candidates per design module (RFantibody, IgGM, and mBER).

This approach allowed for parallel execution of multiple design jobs while maintaining traceability of design parameters. The manifest-driven architecture facilitated efficient GPU allocation and provided a structured framework for tracking and aggregating results from multiple concurrent design runs. Similar batch processing strategies were subsequently employed in the scoring phase, where structural prediction and evaluation tools were executed across the consolidated design pool.

The computational infrastructure consisted of 2,000 GPUs distributed across two AWS regions (1,000 GPUs each in us-east-1 and us-west-2). GPU instance selection was optimized based on regional availability, with us-east-1 primarily utilizing g5 instances and us-west-2 leveraging g6e instances. This multi-region strategy provided robust computational capacity while maintaining flexibility to accommodate varying instance availability patterns.

### 5.3 Antigen structure generation

5 different antigen structures were used in the de novo design. Structure A is retrieved from AlphaFold Protein Structure Database [23]. Structure B,C,D,E are predicted using open-source protein folding models with the same key parameters including recycling steps set to 10, number of diffusion samples set to 15, and multiple sequence alignment (MSA) enabled. Structure B and C are generated by Boltz-2 [7], with structure B enabling argument use_potentials and structure C disabling argument use_potentials. Structure D is generated by Chai-1 [8], and structure E is generated by IntelliFold [9].

### 5.4 Hotspot Recommendation Agent and Tools used

The agent operates through a deterministic workflow that orchestrates seven complementary bioinformatics tools. An example of the agent’s recommendation is shown in Table 1.

**Table 1:**
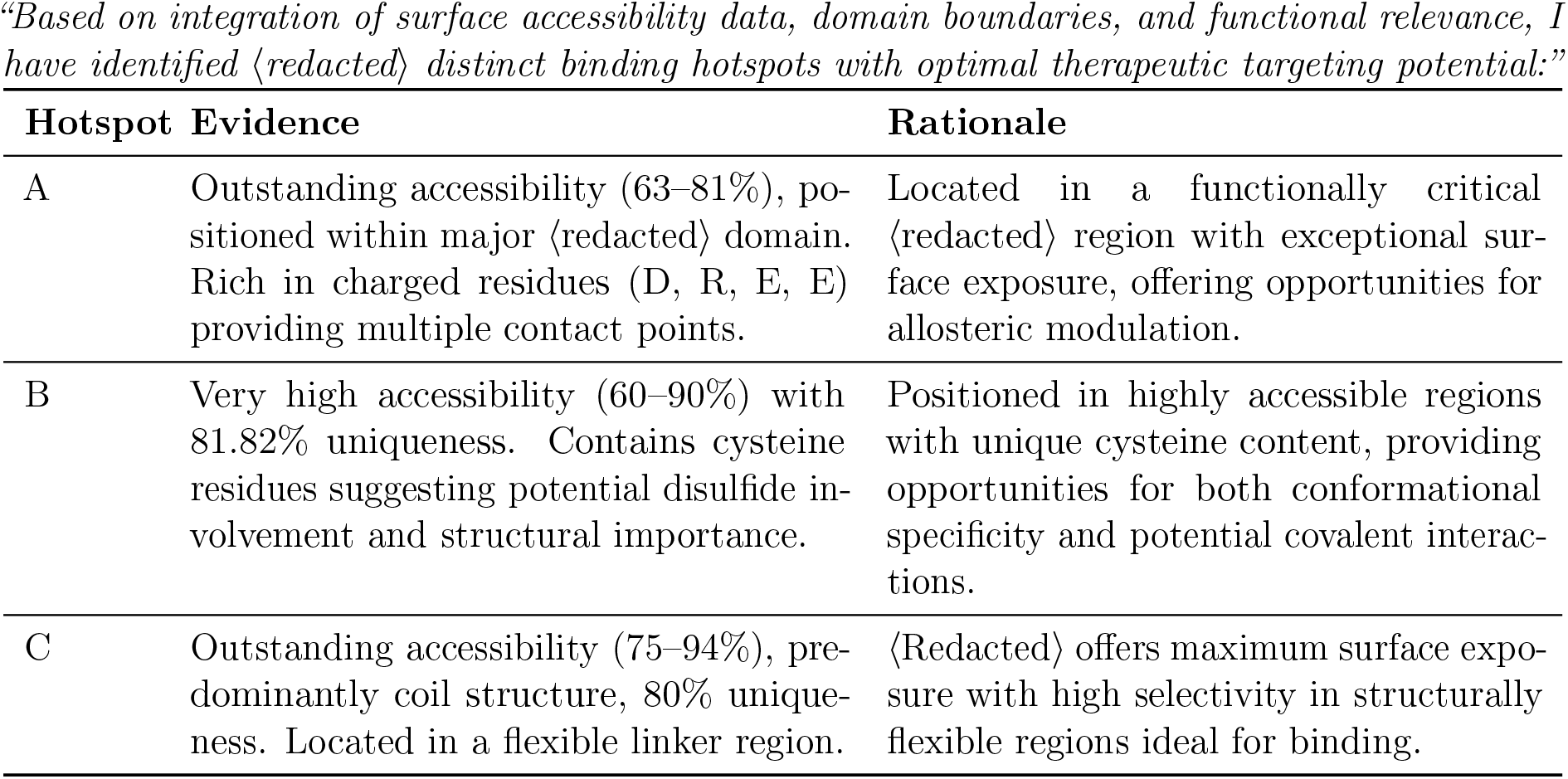
Representative hotspot recommendations from the agent. Three of the eight recommended epitope regions are shown with supporting evidence and rationale. Sensitive protein information is redacted with <redacted>.

#### 5.4.1 Unique Regions Analyzer

The user provides both the target antigen they want to bind to and any negative targets they do not want to bind to. The analyzer will perform sequence alignment over these targets to ensure that the agent recommends only epitopes exclusive to the positive targets.

#### 5.4.2 IEDB Epitope Database Search

We align the input antigen sequence against a curated database of 500k epitopes and literature from the Immune Epitope Database (IEDB) using MMseqs2 with optimized parameters for short epitope matching. Default E-value thresholds were manually tuned to accommodate discontinuous conformational epitopes and linear B-cell epitopes.

#### 5.4.3 Protein Family (PFAM) Domain Annotation

Functional domain boundaries provide critical context for hotspot selection. We replaced the computationally expensive InterPro/HMMer3 pipeline with MMseqs2-based PFAM lookup, reducing latency from >10 minutes to <1 minute while preserving annotation quality.

#### 5.4.4 Solvent-Accessible Surface Area

Surface exposure is a prerequisite for antibody accessibility. We compute per-residue SASA, with careful attention to chain isolation, for complex PDB inputs, existing antibody chains must be removed before calculation to avoid artificially reduced accessibility scores at the true binding interface.

#### 5.4.5 Secondary Structure

Three-state secondary structure classification (helix, sheet, coil) informs epitope likelihood, as loop regions and beta-turns are overrepresented in known epitopes compared to buried helical segments.

#### 5.4.6 Hydrophobicity Profiling

Per-residue hydrophobicity scores characterize the physicochemical landscape of the target surface. Antibody-antigen interfaces typically feature a mixed composition, with hydrophobic residues contributing to binding energy at the interface core and surrounding polar or charged residues providing specificity and solvent-mediated contacts. The profiling identifies both hydrophobic patches that may drive binding affinity and flanking hydrophilic regions that may contribute to epitope accessibility and specificity.

#### 5.4.7 Interface Analyzer

When users provide antibody-antigen complex structures, this tool directly computes interface contacts by computing the pairwise distance across all residue pairs for all chains. For this study, our antigen structure had no prior complex, and therefore this tool was not used by the agent.

#### 5.5 De Novo Antibody Design

We used 3 different protein design methods including RFantibody, IgGM and mBER to design CDR I-III regions of 3 different nanobody frameworks for the target antigen. In addition to diversifying de novo design methods, target hotspots and nanobody frameworks, we designed against 5 different predicted target structures (each from a different structure prediction model), systematically varied CDR III length and ran de novo methods with multiple random seeds to increase the diversity of our nanobody designs. For each de novo method, we generated 96,000 nanobody designs by varying the parameters and designed 288,000 nanobody sequences in total.

#### 5.5.1 RFantibody for de novo antibody design

We used RFantibody, a structure-based generative model that leverages denoising diffusion probabilistic models (DDPMs), to design nanobody CDR backbones on a fixed framework. RFantibody operates by iteratively denoising random 3D coordinates into biophysically plausible backbone structures. For antibody-antigen applications, the model conditions CDR loop design on the fixed antigen structure. The hotspot residue set is an input parameter and is specified as soft constraints that bias the diffusion trajectory toward generating conformations to the intended epitope.

#### 5.5.2 Optimized Implementation for RFantibody

We developed an accelerated implementation of RFantibody that achieved 2 to 7x speedup over the vanilla RFantibody release with minimal performance degradation. Our optimizations target two computational bottlenecks: a) SE(3) Transformer Quantization. We identified layers within the SE(3)-equivariant transformer architecture that exhibit low sensitivity to reduced numerical precision. The weight layers and input/output activations inside the transformer were selectively quantized to BF16/FP16, substantially reducing memory bandwidth requirements and enabling increased batch throughput without measurable impact on design quality metrics (interface RMSD, pLDDT, PAE). b) IGSO(3) Distribution Caching. The diffusion process over rotational degrees of freedom requires sampling from the IGSO(3) distribution, the isotropic Gaussian distribution on SO(3). Computing these distributions at runtime introduces significant latency. We precompute and cache IGSO(3) probability densities across the relevant parameter ranges, eliminating redundant computation during inference.

To assess the quality of the optimized implementation, we performed two levels of validation. First, we measured the per-layer quantization error ∥*W*_*q*_ · *a*_*q*_ − *W* · *a*∥ between the quantized weights *W*_*q*_ and original weights *W* applied to representative activations *a* from the previous layer, and confirmed that the error remained within the 16-bit machine epsilon bound, consistent with the theoretical precision floor of 16-bit reduced-mantissa formats. Second, we evaluated design quality end-to-end by running both the original and optimized RFantibody on generated samples across a diverse set of therapeutic targets and nanobody frameworks. The designed sequences from each implementation were independently folded using RF2 to recover backbone structures. The distributions of interaction PAE, predicted LDDT, and framework-aligned CDR RMSDs were near-equivalent between the two implementations, confirming that the 2–7x speedup does not affect design quality.

#### 5.5.3 IgGM for de novo antibody design

We used IgGM, a structure-conditioned deep generative model, to design nanobody CDR sequences on a fixed framework given the predicted 3D structure of the target antigen. IgGM uses the diffusion process to jointly optimize CDR regions on nanobody sequences and structure of the nanobody-antigen complex. It takes nanobody framework with CDR residues masked and predicted antigen structure as inputs and outputs a nanobody sequence with filled-in CDR I-III regions. When designing with IgGM, we create a design run to generate 100 nanobody sequences, each with a random seed for the diffusion process. We complete a total of 960 runs, where each run is parametrized by 3 different nanobody frameworks, 5 predicted antigen structures, 8 different CDR III lengths (6 - 13 residues) and 8 target hotspots (3 x 5 x 8 x 8 = 960). We generate a total of 96,000 nanobody designs with IgGM.

#### 5.5.4 mBER for de novo antibody design

We used mBER, a structure- and sequence-conditioned binder-design framework, to design nanobody CDRs on a fixed VHH scaffold given the 3D structure of the target antigen. mBER leverages a fixed nanobody framework, masks the CDR regions, and then uses gradient-based optimization of sequence logits through a structure prediction model (AlphaFold-Multimer), guided by sequence priors from a protein language model, to iteratively optimize the variable regions for binding. Rather than updating model weights, mBER backpropagates structure prediction losses to the input sequence representation to identify CDR sequences predicted to form stable bound complexes. The method outputs fully specified nanobody sequences (within the defined framework) that are predicted to adopt stable bound structures against the target antigen epitope. Similar to IgGM, we use the same parameters to complete 960 design runs and generate a total of 96,000 nanobody designs with mBER.

### 5.6 Antibody Scoring Workflow

Prior to implementing the scoring phase, we consolidated all nanobody designs from multiple design methods (RFantibody, IgGM, and mBER) into a unified dataset. This aggregation step involved collecting designs generated against different predicted target structures, various hotspots, and across different nanobody frameworks. Each design entry was standardized to include the full nanobody sequence, design method parameters, and target epitope information to facilitate subsequent systematic evaluation. This consolidated dataset of 288,000 initial candidates was then prepared for comprehensive scoring using multiple structural prediction tools including NanobodyBuilder2 and Boltz-2.

For optimal computational resource utilization, we implemented a partitioned batch processing strategy. The 288,000 candidates were divided into smaller batches optimized for concurrent execution across the available GPU resources. Within each batch, sequences were processed sequentially to maintain computational stability. This batching approach balanced the need for high-throughput processing with the constraints of limited GPU availability, enabling efficient scoring of the large candidate pool.

#### 5.6.1 NanoBodyBuilder2 Antibody Folding

For structural validation of the nanobody designs, we employed NanobodyBuilder2 to evaluate the structural integrity of individual candidates. Prior to scoring, all design candidates were consolidated into a single dataset. Each nanobody sequence was processed through NanobodyBuilder2, which generates predicted structural metrics including pLDDT scores. These scores serve as a proxy for structural plausibility and stability of the designed nanobodies in their unbound state. This structural validation step formed one component of our multi-metric scoring approach used to evaluate and filter candidate designs.

#### 5.6.2 Boltz-2 Antigen-Antibody Co-Folding

Complex structure prediction and validation were performed using Boltz-2. For each nanobody-antigen pair, Boltz-2 was used to predict the structure of the complex, generating key metrics including complex iPTM and complex ipLDDT scores. These metrics served as indicators of binding plausibility and complex stability.

Additionally, from the predicted complex structures, we calculated minimum CDR distances by identifying the complementarity-determining regions (CDRs) using Abnumber and measuring the minimum atomic distances between CDR residues and the antigen.

#### 5.6.3 Parameters Used (Default Settings)

The following parameters were configured for the Boltz-2 prediction:

`recycling_steps`**: 3** The number of recycling steps to use for prediction.

`sampling_steps`**: 200** The number of sampling steps to use for prediction.

`diffusion_samples`**: 1** The number of diffusion samples to use for prediction.

`max_parallel_samples`**: 5** Maximum number of samples to predict in parallel.

`step_scale`**: 1.638** The step size is related to the temperature at which the diffusion process samples the distribution. The lower the value, the higher the diversity among samples (recommended between 1 and 2).

#### 5.6.4 MochiBind Antigen-Antibody Binding Affinity Prediction

MochiBind [21] is our internally developed sequence-based binding affinity predictor to estimate binding affinities of our designs to the target antigen to get a final scoring metric that’s orthogonal to structural metrics discussed earlier. MochiBind uses ESM2 to embed the nanobody-antigen complex and applies a predictor head on the complex embedding to predict the binding affinity of the nanobody sequence. Our validation results indicate that this method outperforms structurebased proxies for binding affinity on out-of-distribution datasets of unseen antigen-antibody pairs. Using MochiBind at the scoring stage, we predicted the binding affinity of each nanobody design against our target antigen and used this metric for downstream filtering. MochiBind manuscript is currently under review.

#### 5.6.5 Antibody Diversity and Liability Analysis

Sequence diversity module employed IMGT-numbered positional analysis to determine amino acid frequencies across Complementarity Determining Regions (CDRs) and Framework Regions (FRs). Sequence-based clustering was performed using MMseqs2 for enhanced performance and accuracy, complemented by traditional metrics like substitution analysis and pairwise distance calculations to quantify sequence variability. The liability analysis module systematically checked for a range of potential developability risks, adopting checks from the LAP methodology [29]. This check was designed to look for critical sequence motifs for high-risk liabilities such as N-linked glycosylation, high-risk deamidation, and fragmentation within CDRs and FRs, along with other concerns like isomerization, cysteine bond issues, and oxidation risks, providing an essential assessment of potential manufacturing and stability challenges. Antibody candidates which have red flags from the liability check were filtered out.

### 5.7 Candidate Selection

Computationally designed nanobody candidates must be evaluated across multiple competing structural and functional quality metrics before advancing to experimental validation. Selecting the top 100,000 candidates from a large, filtered library requires balancing these objectives without imposing arbitrary relative weights, preserving diverse high-quality candidates rather than collapsing to a single composite score. The 288,000 de novo designed VHH candidate sequences as well as metrics from the scoring modules were selected through multi-property optimization through Pareto ranks to select the 100,000 best candidates that perform well in five objectives shown in Table 2.

**Table 2:** Properties used for antibody candidate selection.

| Property | Description |
| --- | --- |
| CDR Min Distance | Minimum distance between CDR loops and the target, inverted so that tighter engagement scores higher |
| Predicted Binding Affinity | MochiBind predicted binding affinity (inverted), reflecting stronger target interaction |
| Protein iPTM | Interface predicted TM-score from structure prediction, measuring confidence in the binding interface |
| Complex ipLDDT | Interface per-residue local distance difference test, assessing local structural accuracy at the interface |
| Nanobody Avg pLDDT | Average pLDDT across the nanobody, reflecting overall fold confidence |

Each candidate was scored on five normalized properties (higher is better):

We applied a modified version of the Efficient Non-dominated Sort (ENS) algorithm to identify non-dominated antibodies to stratify all candidates into successive Pareto fronts without requiring objective weights:

- Pareto Front Identification. A candidate belongs to Rank 0 (the true Pareto front) if no other candidate is simultaneously equal or better on all five objectives and strictly better on at least one. This front represents the set of candidates where improving any single metric necessarily comes at the cost of another.
- Iterative Peeling. After removing Rank 0, the same non-dominated sorting is repeated on the remaining candidates to identify Rank 1, then Rank 2, and so on. This produces an ordered layering of the full library by multi-objective quality.
- Pareto Rank Thresholding. We computed the minimum number of Pareto ranks that resulted in our target library size of 100,000. Specifically, we selected the first 9 Pareto ranks completely and randomly sampled from the Pareto Rank 10 set until we had the required number of antibodies.

### 5.8 Yeast Library Construction and Quality Control

Linear DNA libraries were synthesized by Twist Bioscience and transformed into the *Saccharomyces cerevisiae* strain EBY100. Library quality control was performed using both Sanger sequencing and Next-Generation Sequencing (NGS) with the MiSeq 300×300 kit (Illumina), adhering to the manufacturer’s recommended protocol.

### 5.9 Yeast Display Induction and Staining

Transformed yeast cells were cultured in SDCAA medium (Teknova) and subsequently induced at an OD_600_ of 1.0 for 48 hours at 18°C in SGCAA medium (Teknova). Following induction, cells were washed with PBS supplemented with 0.5% BSA (PBSA). The cells were incubated with the target antigen for 1 hour at 4°C with gentle shaking. Post-incubation, cells were centrifuged at 300×g for 5 minutes and washed with PBSA.

For fluorescent labeling, cells were resuspended in 1 mL of buffer containing secondary antibodies: 1:200 of 0.5 mg/mL anti-Flag-647 (GenScript) for antigen detection and 1:200 of 0.5 mg/mL anti-V5-488 (GenScript) for VHH display quantification. The mixture was incubated for 1 hour at 4°C. Cells were washed three times with PBSA and finally resuspended in 1 mL of PBSA for analysis.

### 5.10 Flow Cytometry and Sorting

Sorting was performed using an SH800S Cell Sorter (Sony Biotechnology). The gating strategy isolated the double-positive population (indicating both VHH display and antigen binding) as well as the display-positive/antigen-negative population. A minimum of 10-fold coverage of the library diversity was bulk sorted and propagated for the subsequent round of selection.

In Round 1, cells were sorted against 400 nM target antigen. In Round 2, high non-specific background binding to the secondary detection antibodies was observed. To mitigate this, a negative sorting step was introduced prior to positive selection: the induced library was incubated with all secondary detection reagents without antigen, and the non-binding population was collected to deplete clones exhibiting non-specific adhesion. Positive sorting was then performed at 200 nM target antigen, with both higher and lower binding subpopulations within Q2 collected separately.

### 5.11 SPR

The 116 selected VHH candidates were recombinantly expressed in HEK293 cells as VHH-fc fusions and purified via protein A chromatography. All 116 candidates expressed successfully, with a median yield of 368 *µ*g per 2 × 1 mL culture (range: 69–400 *µ*g; 97% above 200 *µ*g). Binding kinetics were characterized by surface plasmon resonance (SPR) on a Carterra LSA instrument. The VHH-fc was immobilized on the sensor surface, and the target antigen ECD were injected as analytes at multiple concentrations to determine association rate (*k*_*on*_), dissociation rate (*k*_*off*_), and equilibrium dissociation constant (*K*_*D*_). Specificity was confirmed by testing all candidates against an irrelevant control antigen transferrin receptor protein (TfR1); no binding was detected for any candidate against the control, confirming target-specific interaction. Candidates with measurable binding responses and *R*_max_ ≥ 30 RU were considered to have reliable kinetic fits for affinity determination.

## 6 Acknowledgement

The authors acknowledge the AWS Life Sciences team and the AWS HealthOmics team for providing resources and input that contributed to the results reported within this paper. We also thank Brian Loyal from AWS HCLS for reviewing the manuscript.

## Author Contributions

Y.Z., M.Y., E.L., C.T., L.G., K.S., J.K., X.C. conceived and designed the work. C.T., M.Y., E.L., Y.Z. implemented the de novo design and scoring workflow. K.S., Y.Z., M.Y., C.T., E.L. performed the candidate selection. M.E.C. and N.K.V.C. discovered the tumor target through omics carried out at MSK. J.K., X.C., M.E.C., and N.K.V.C. supervised the project. All authors discussed the results and approved the final version of the manuscript.

## Supplementary Information

**Table S1:** Pairwise structural similarity (TM-score) of predicted models for the target protein.

|  | AlphaFold2 | Boltz-2 | Boltz-2 + potential | Chai-1 | IntelliFold |
| --- | --- | --- | --- | --- | --- |
| AlphaFold2 | — | 0.627 | 0.632 | 0.663 | 0.622 |
| Boltz-2 |  | — | 0.909 | 0.850 | 0.775 |
| Boltz-2 + potential |  |  | — | 0.864 | 0.825 |
| Chai-1 |  |  |  | — | 0.893 |
| IntelliFold |  |  |  |  | — |

TM-scores were computed on C $\alpha$ atoms after optimal superposition. A TM-score > 0.5 indicates the same global fold; 1.0 denotes identical structures.

**Figure S1:**
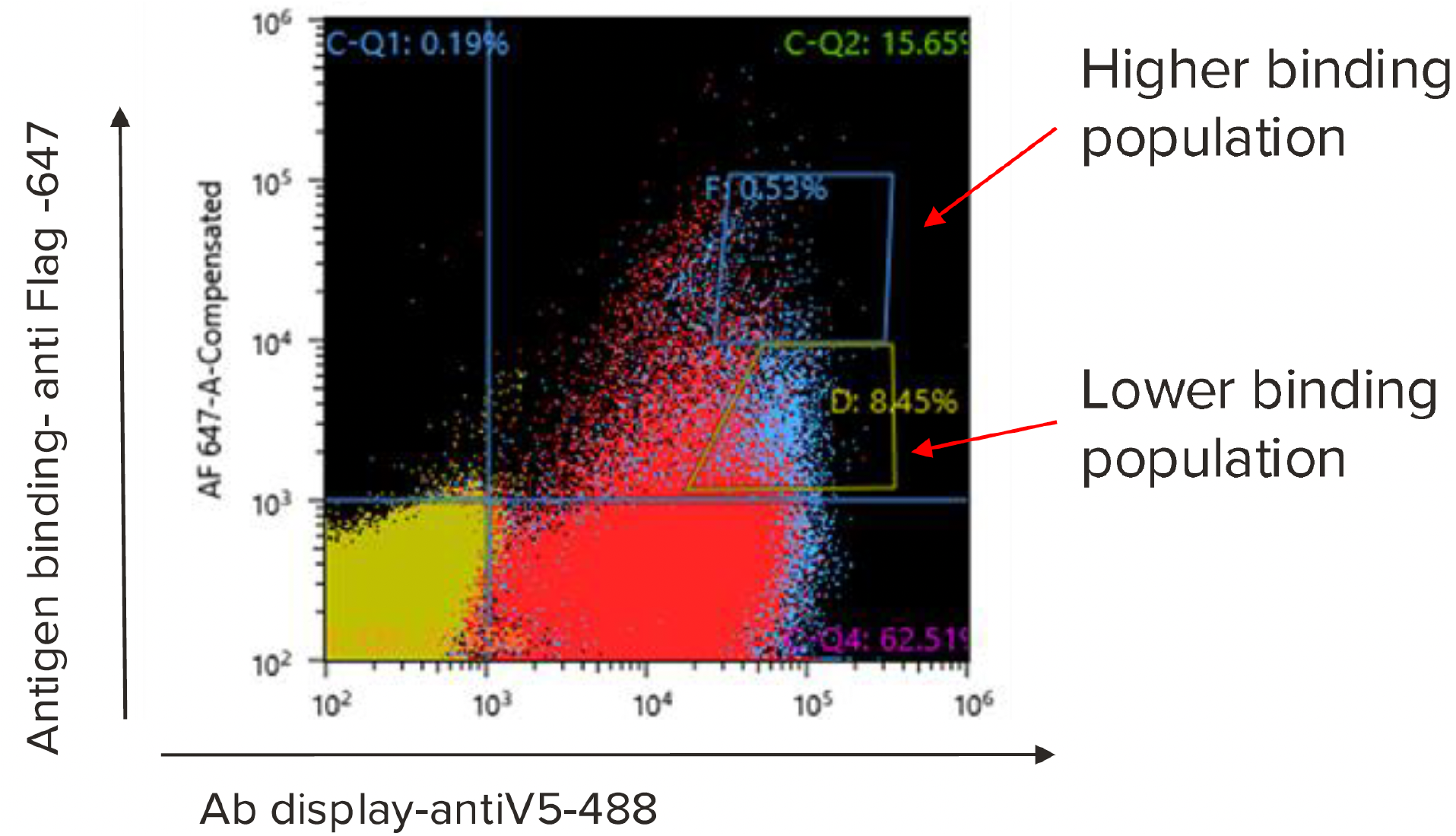
Round 2 FACS enrichment and selection of 116 candidates for SPR. Following the 2nd round of FACS enrichment, individual yeast clones from the binding-positive population were arrayed and rescreened by flow cytometry. Binding specificity for each clone was assessed using mean fluorescence intensity (MFI) measurements. Clones exceeding a predefined MFI threshold were classified as specific binders, yielding 116 candidates that were advanced to recombinant expression, purification, and SPR characterization.

**Figure S2:**
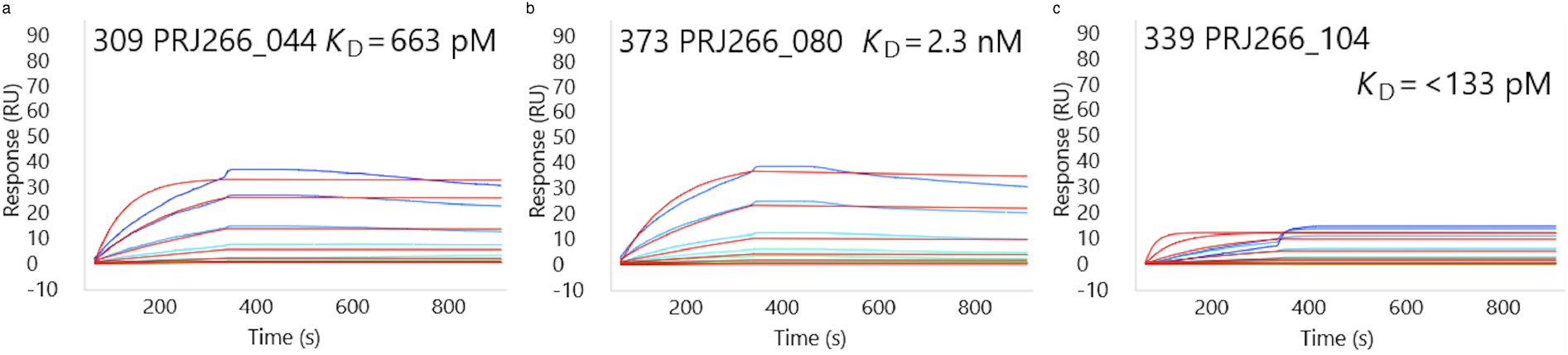
SPR sensorgrams of three top de novo designed VHH binders against the target antigen. Multi-concentration kinetic binding curves showing association and dissociation phases for (a) PRJ266_044, (b) PRJ266_080, and (c) PRJ266_104. Colored traces represent experimental data at different analyte concentrations; fitted curves are overlaid.

**Figure S3:**
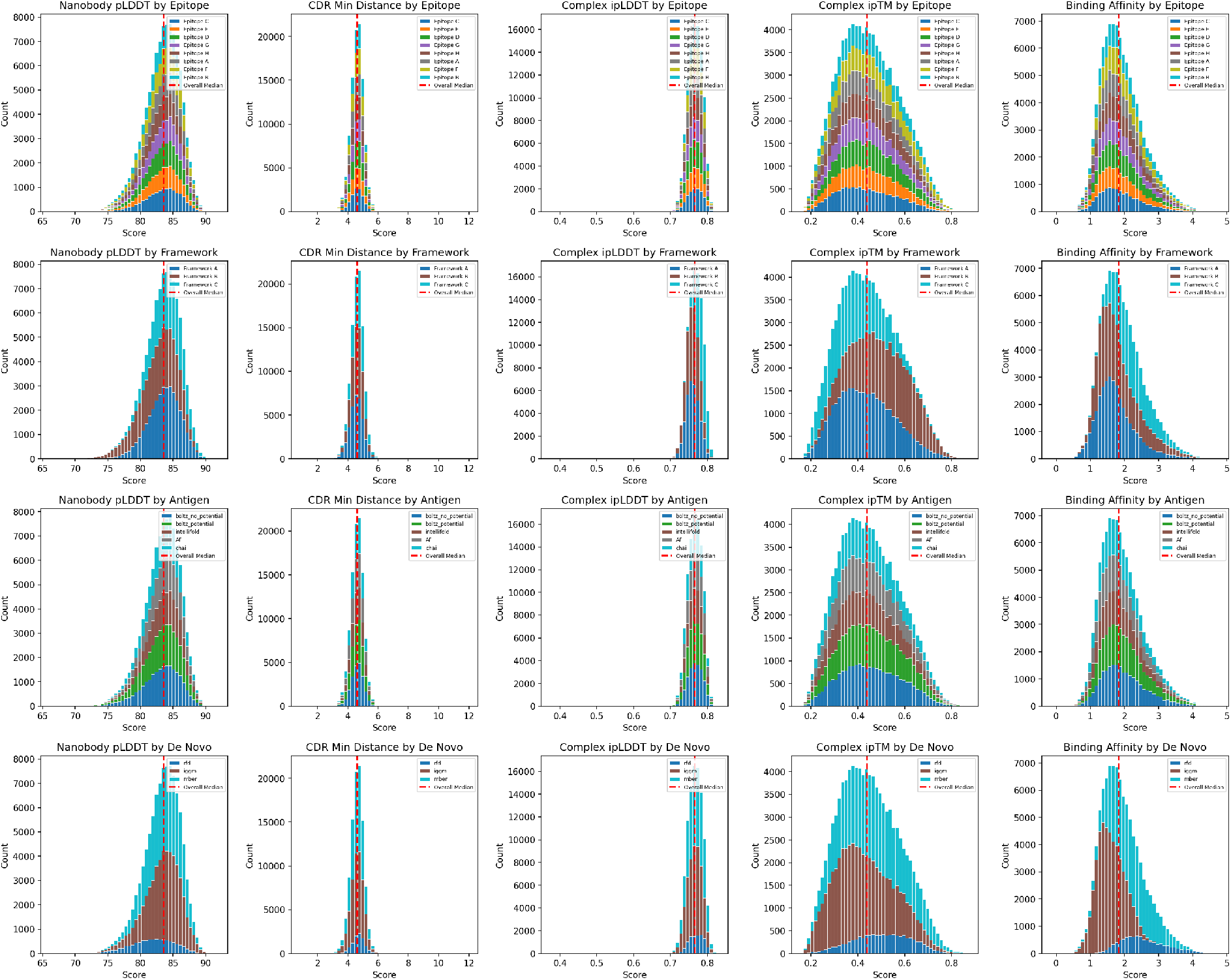
Distribution of *in silico* metrics for 100,000 candidates selected for *in vitro* experiments. Rows correspond to experiment parameters (from top to bottom): target epitope, nanobody framework, antigen structure prediction method, de novo design method. Columns correspond to *in silico* metrics (left to right): nanobody monomer folding pLDDT score, minimum CDR-antigen distance, complex ipLDDT, ipTM and predicted binding affinity. Higher scores indicate better designs, except in minimum CDR distance and predicted binding.

**Figure S4:**
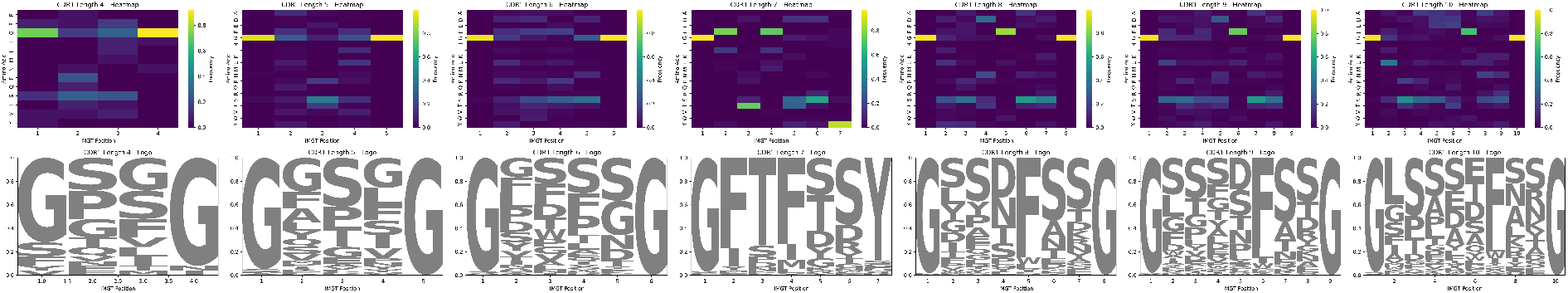
Library sequence diversity analysis for CDR H1 region positional amino acid frequencies for different CDR lengths. Heatmaps and sequence logos visualize the same underlying frequencies for a given CDR length.

**Figure S5:**
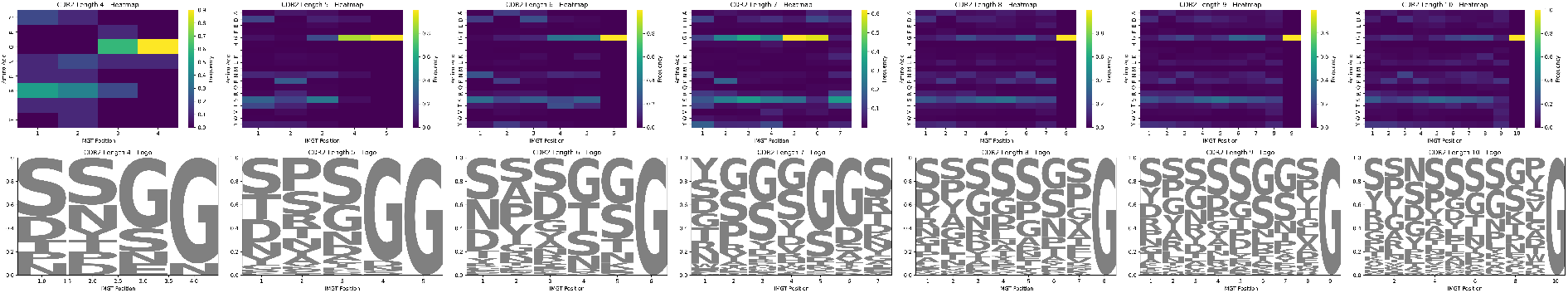
Library sequence diversity analysis for CDR H2 region positional amino acid frequencies for different CDR lengths. Heatmaps and sequence logos visualize the same underlying frequencies for a given CDR length.

**Figure S6:**
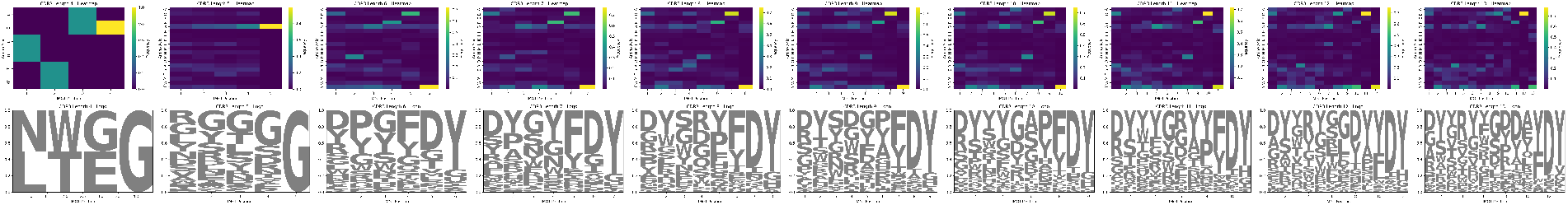
Library sequence diversity analysis for CDR H3 region positional amino acid frequencies for different CDR lengths. Heatmaps and sequence logos visualize the same underlying frequencies for a given CDR length.

**Figure S7:**
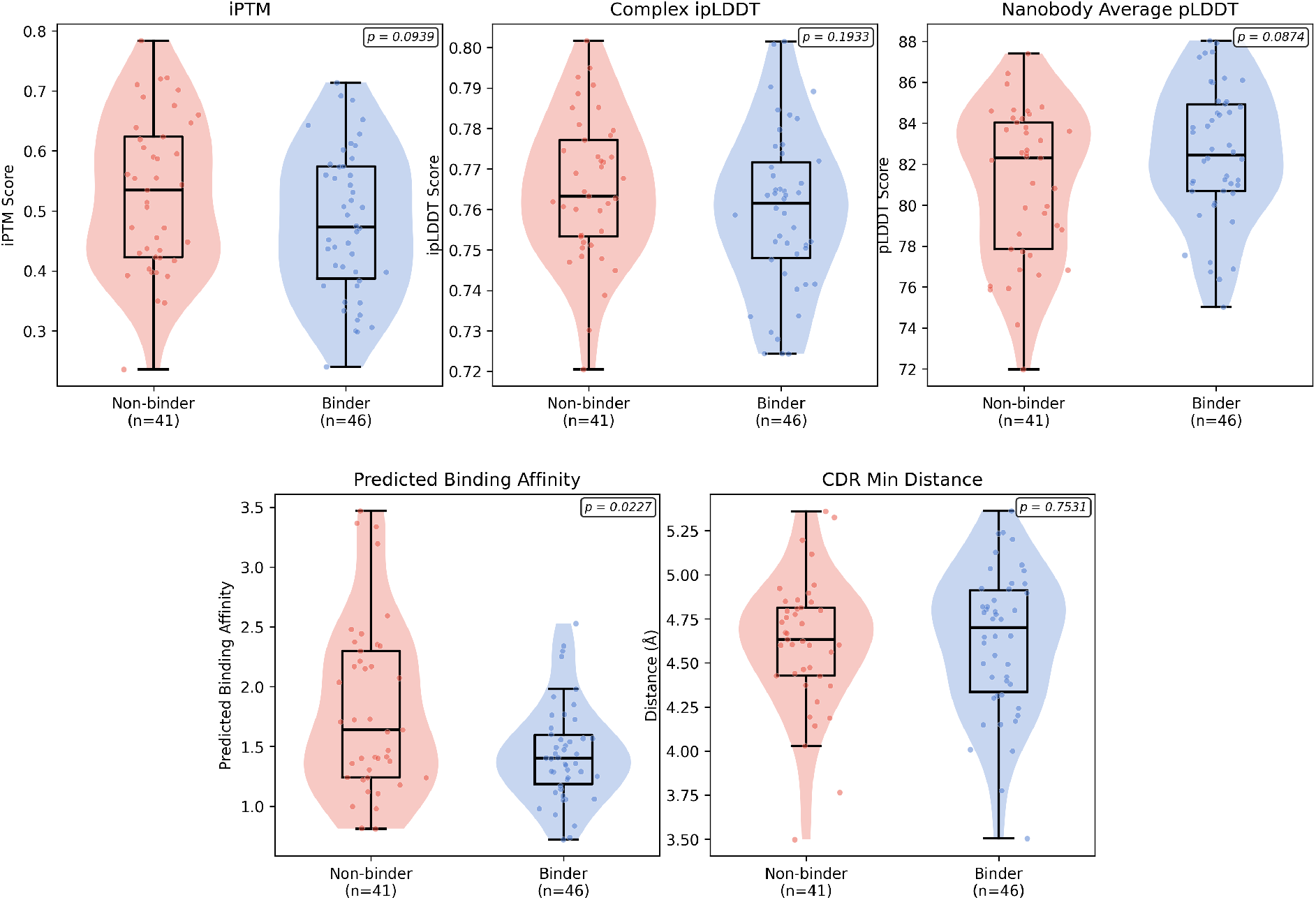
Comparison of *in silico* metrics for SPR-confirmed binders and non-binders. Violin plots comparing five computational metrics between nanobodies classified as binders (*R*_max_ ≥ 30 RU, n=46) and non-binders (n=41) from the 116 SPR-characterized candidates. Features include Boltz-2 interface predicted TM score (iPTM), complex interface predicted LDDT (ipLDDT), NanoBodyBuilder2 average pLDDT, MochiBind-predicted binding affinity, and minimum CDR-antigen distance. Box plots show median and interquartile range, p-values from two-sided Mann-Whitney U tests. MochiBind-predicted binding affinity shows the strongest discrimination between groups (p = 0.0227).

**Figure S8:**
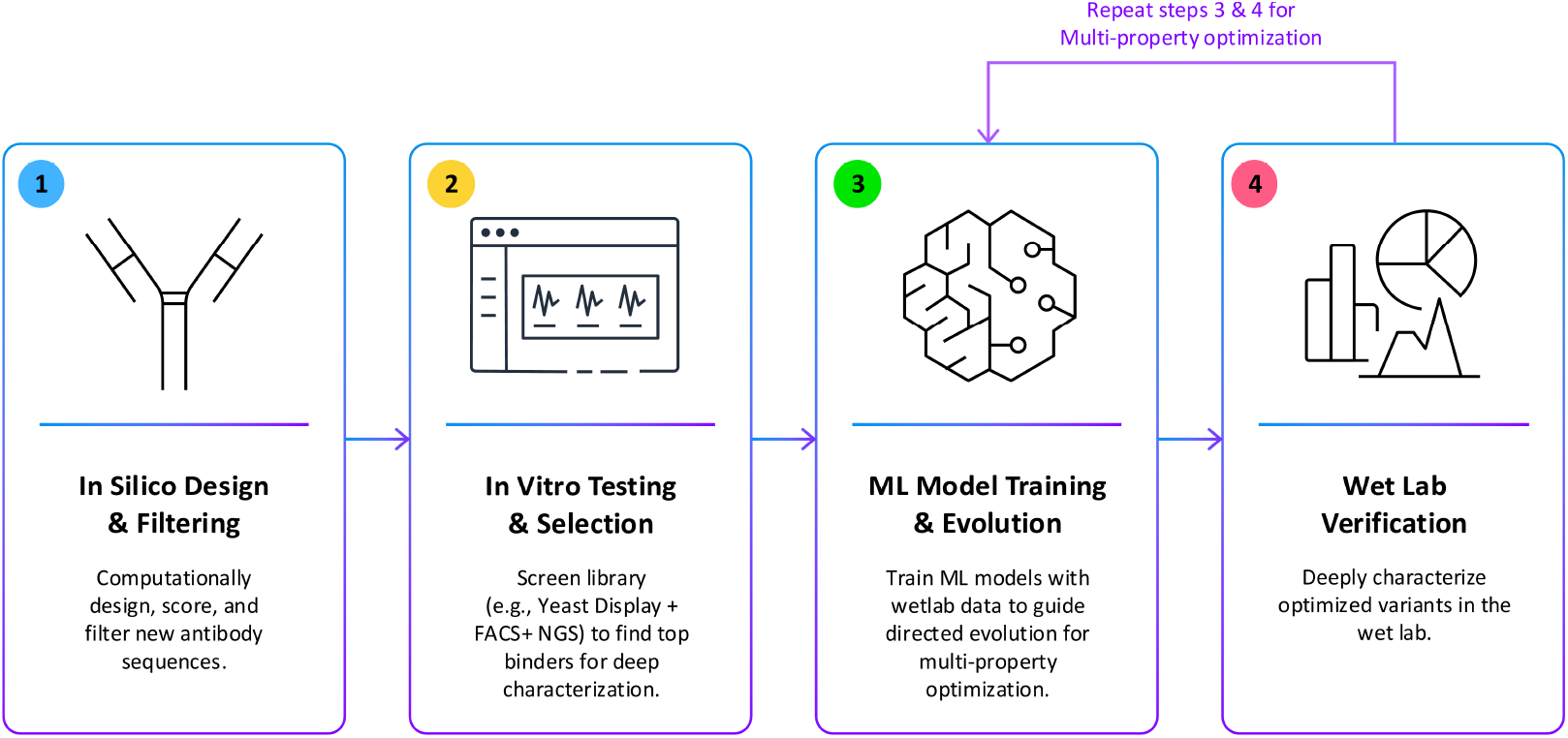
Design-Build-Test-Learn (DBTL) cycle enabling iterative multi-property optimization through Lab-in-the-Loop integration. The workflow comprises four interconnected stages: (1) *In Silico* Design & Filtering (blue) computationally generates, scores, and filters new antibody sequences using de novo design methods and multimetric evaluation. (2) *In Vitro* Testing & Selection (yellow) screens libraries using high-throughput experimental methods (yeast display, FACS, NGS) to identify top binders for deep characterization. (3) ML Model Training & Evolution (green) trains target-specific machine learning models on experimental data to predict binding affinity, stability, and developability, guiding directed evolution campaigns. (4) Wet-lab Verification (red) performs comprehensive biophysical characterization of optimized variants including binding kinetics, thermal stability, and other therapeutic properties. The cycle iterates through stages 3 and 4, progressively refining candidates through ML-guided evolution and experimental validation. This closed-loop framework couples computational predictions with experimental feedback to systematically improve antibody properties while maintaining a mechanistic understanding of design-function relationships.

